# Ancestral Sequences Cannot be Accurately Reconstructed via Interpolation in a Variational Autoencoder’s Latent Space

**DOI:** 10.1101/2025.11.19.689264

**Authors:** Evan Gorstein, Mengze Tang, Hailey Bruzzone, Claudia Solís-Lemus

## Abstract

Standard methods for ancestral sequence reconstruction (ASR) rely on substitution models for the residues in a biological sequence and assume independent evolution across these sites, ignoring the epistatic interactions that shape molecular evolution. In contrast, deep learning models like variational autoencoders (VAEs) can learn low-dimensional representations (“embeddings”) of sequences in a protein family that may implicitly handle these dependencies, raising the possibility of performing more accurate ASR by interpolating between extant sequence embeddings within the VAE’s latent space. In this study, we test this hypothesis by developing and evaluating a VAE-based ASR pipeline. Benchmarking this approach against established likelihood-based and parsimony methods using various simulations of protein evolution, including scenarios with and without epistasis, we find that the VAE-based approach is consistently and significantly outperformed by standard methods, even in epistatic regimes where it was hypothesized to have an advantage. We further show that this failure is not due to a lack of phylogenetic structure in the latent space, which does contain evolutionary signal. Rather, the primary limitation is the information loss inherent to the autoencoding process: the VAE’s decoder cannot generate sequences with sufficient fidelity for the precise demands of ASR.

## 1 Introduction

Substitution models which characterize how individual sites in a biological sequence evolve stochastically over time have served as a workhorse in molecular evolutionary research, facilitating the study of the evolutionary history of a gene or protein family, including the estimation of the relative timing of residue substitutions, gene duplications, and speciation events (Jukes et al. 1969; Dayhoff et al. 1978; Whelan and Goldman 2001; Le and Gascuel 2008). Application of these models consists in viewing the sequence as a collection of independently evolving residues, which in turn allows one to avoid the combinatorial explosion of sequence space when computing the likelihood of a given evolutionary scenario. The resulting computational tractability comes at the cost of neglecting the epistasis that often plays a role in molecular evolution, which can occur due to interactions between sites in determining structure and function of a protein (Starr and Thornton 2016; Nasrallah et al. 2010).

Outside of phylogenetics, researchers have increasingly leveraged methods from machine learning and natural language processing to study protein sequences. In particular, there has been much work in recent years towards the development of unsupervised or weakly supervised methods to learn useful representations of protein sequences as dense numerical vectors, known as “protein embeddings”, from the large collections of homologous protein sequences deposited and cataloged in online databases (Yang et al. 2018; Rao et al. 2019; Detlefsen et al. 2022). Although these embeddings have more recently become associated with large protein language models trained on millions of sequences from thousands of diverse families (Fan et al. 2025), prior to the proliferation of these universally trained models, researchers trained models such as variational autoencoders (VAEs) to learn embeddings for only the sequences in an individual protein family. The resulting embeddings have been used to visualize the space of sequences within the family, predict the impact of mutations, and even generate novel viable sequences (Sinai et al. 2017; Riesselman et al. 2018; Hawkins-Hooker et al. 2021; Ziegler et al. 2023). Detlefsen et al. (2022) discuss the various choices involved in learning protein representations and illustrate empirically the implications of these choices for downstream predictive and interpretive goals. Notably, they show that when the goal is to represent the phylogenetic relationships within a given family, training on aligned sequences from that family alone is advantageous, although state-of-the-art protein language models trained universally do also learn within-family phylogenetics (Tule et al. 2024).

Exploring the shape of the manifold in latent space of the embeddings from a VAE trained on a single family of aligned sequences, both Detlefsen et al. (2022) and Ding et al. (2019) find that it captures the evolutionary structure of the data, namely, that sequences have evolved along a phylogenetic tree. In the latter article, the authors hierarchically cluster the embeddings based on Euclidean distance within the latent space and find that these clusters identify larger evolutionary clades more accurately than the clades in a tree estimated with FastTree (Price et al. 2009); however, these embedding clusters are less accurate than the estimated tree at identifying smaller clades. In theory, the advantage of embedding models relative to the independent-sites models employed by phylogeneticists is their flexibility, model-agnosticism, and potential to capture epistatic effects by learning directly from the data. The results from Ding et al. (2019) suggest, however, that although the global organization of the embeddings recapitulates the overall phylogeny, finer-grained historical relationships may not be recoverable from them.

In this paper, we investigate the limits of an embeddings-based approach to phylo-genetics by studying whether we can perform ancestral sequence reconstruction (ASR) by interpolating within the latent space of a VAE. Ancestral sequence reconstruction is a fundamental task in phylogenetics in which the analyst attempts to infer, on the basis of extant homologous sequences, the amino acids of an ancestral sequence at one or multiple of the internal nodes of the phylogenetic tree that relates these homologs (Pauling et al. 1963; Hochberg and Thornton 2017; Selberg et al. 2021). Classically, ancestral sequence reconstruction was done using a maximum parsimony principle (Dayhoff et al. 1978), but likelihood-based approaches have emerged as the dominant methodology in more recent years. The models that these approaches assume and from which the likelihood is thus derived are the substitution models mentioned at the out-set, along with the corresponding implicit assumption of independent evolution across sites. In this paper, we benchmark an approach to ASR based on the embeddings from a VAE, comparing it to this more standard approach based on the likelihood function from substitution models. In addition to learning a function for embedding sequences in a continuous space (the encoder), VAEs simultaneously learn a mapping back to the space of sequences (the decoder) that is trained to approximate the inverse of the encoder. Thus, interpolation within the embedding space of VAE can be used to generate novel, yet meaningful data by feeding the interpolates through the decoder. In the context of ASR, the novel, meaningful data we want our variational autoencoder to produce are the ancestral sequences that once existed in the past and which gave rise to the extant sequences on which the VAE is trained.

Following a brief review of related work, we detail this approach, scalable to families with thousands of sequences, of embedding-based ancestral sequence reconstruction. We then describe the results of simulation studies conducted to benchmark this approach against existing methods for ASR. In the absence of recoverable ancient sequences, real protein families lack a definitive ground truth with which to evaluate proposed ancestral sequences and it is therefore common to benchmark ASR on computationally simulated evolution (Williams et al. 2006; Moreta et al. 2022; De Leonardis et al. 2025). In this work, we study several different sequence evolution simulation schemes, including multiple that incorporate epistasis or co-evolution of sites, to see whether an embeddings-based approach to ASR might improve the reconstruction accuracy relative to existing methods that ignore epistasis. We find that in each tested scenario, embedding-based ASR is outperformed by standard likelihood-based methods, even when the assumption of independent site evolution made by these methods is violated. In order to do ASR with embeddings, one requires both 1) a latent space that is phylogenetically meaningful and 2) a mapping from sequence to embedding space that does not lose too much information about the sequence. In the context of the simulations with epistasis, we discuss the relative responsibility of each condition’s absence for the inferior performance of an embeddings-based approach to ASR.

## 2 Related work

While there has been effort in the phylogenetic community to develop models of protein sequence evolution that incorporate epistatic effects, with the notable exception of the recent work of De Leonardis et al. (2025), there are few methods for performing phylogenetic inference based on these models. Robinson (2003) and Rodrigue et al. (2006) both consider epistatic models of sequence evolution based on a structure-based fitness landscape, but these models have not been used for phylogenetic inference. Meyer et al. (2019) models co-evolving sites in sequences of nucleotides in a Bayesian model that facilitates joint inference of the phylogeny and the pairs of coupled sites. De La Paz et al. (2020) propose transforming a Potts model into a dynamic model of sequence evolution with MCMC sampling, which they call Sequence Evolution with Epistatic Contributions (SEEC). While forward simulation from an extension of this model has proven useful for exploring sequence space and generating functional phenotypes (Bisardi et al. 2021; Alvarez et al. 2024), it has not been used for phylogenetic inference.

Recently, De Leonardis et al. (2025) developed an analytically tractable model of sequence evolution that accounts for epistasis and which can be used to perform ancestral sequence reconstruction. In particular, these authors formulate a model according to which each site in a sequences evolves in the context of preceding sites (in some, pre-specified ordering) according to a Markov chain whose stationary distribution is given by the fitted conditional probabilities from a regression on the previous sites. Maximum likelihood ancestral sequence can be reconstructed under this model by following the same ordering, one site at a time, and accounting for the conditional probabilities in reconstructing each position; the resulting estimates are more accurate than reconstructions from IQTree (Minh et al. 2020) in settings where the sequences have been simulated with epistasis.

Finally, Moreta et al. (2022) propose performing ancestral sequence reconstruction with a VAE. While an important motivation for this work, their approach incorporates the known phylogenetic relationships between sequences into the training of the VAE; in particular, they impose a prior on the latent variables in the VAE based on a Gaussian process along the tree, thus tying together the different sequences in the training data. In contrast, we investigate whether an embedding model trained without any access to the phylogenetic tree still learns representations that can be used for accurate ancestral sequence reconstruction, a question that is of interest for several reasons. First, because of the phylogenetic prior in the latent space, the VAE from Moreta et al. (2022) must be trained with full-batch gradient descent, limiting the size of the protein families to which this method can be applied; incorporating the phylogenetic correlation structure only at the later stage in the ASR pipeline via the Brownian motion model applied to pre-computed embeddings enables scaling to larger families with thousands of sequences, as the VAE can be trained with standard mini-batch stochastic gradient descent. Secondly, protein language models are typically trained in a completely unsupervised fashion without phylogenetic tree information available at training time. Excluding information on the phylogenetic tree from the training of our VAE therefore increases the chances that our findings generalize to other attempts at ancestral sequence reconstruction on the basis of protein embeddings. Finally, it has already been demonstrated in Ding et al. (2019) and Detlefsen et al. (2022) that the embeddings from a standard VAE trained on sequences from a large multiple sequence alignment recapitulate phylogeny, so it would not seem necessary to add a phylogenetic prior to the VAE. In section 4.1.1, we reproduce this finding, and in section 4.2, we provide evidence that it is not an absence of phylogenetic signal in the learned embeddings that is causing the poor performance of the embeddings-based approach to ASR that we encounter.

## 3 VAE-assisted ancestral sequence reconstruction

The goal of ASR is to estimate the sequence identity of some determinate ancestor (RNA, DNA, or protein) on the basis of extant homologs from the family. In contrast to other methods for ASR, the embeddings-based approach requires a family of (at least) hundreds of extant sequences, as these comprise the data for training a variational autoencoder. Although in practice the alignment of such a large collection of potentially distantly related sequences and the phylogenetic tree that relates them are both subject to error, especially when estimated based on a potentially flawed model of independent site evolution, for the purpose of focusing solely on evaluating the task of ASR and in line with other ASR methods (De Leonardis et al. 2025; Moreta et al. 2022), we assume that we are in possession of the ground-truth alignment and the ground truth phylogenetic tree, with branch lengths, possibly unrooted. Note as well that if the extant sequences in the alignment are not all orthologs, but also include paralogs, the tree will include some internal nodes representing duplication, rather than speciation, events. Most methods for performing ASR, including the approach presented here, are agnostic to the interpretation of the internal nodes of the tree.

In the pipeline under study, represented graphically in Figure 1, we begin by training a variational autoencoder on the collection of fixed length extant sequences from the alignment to learn a latent variable model of the distribution of sequences in the family. The variational autoencoder framework includes an encoder, which accepts as input a sequence and outputs the parameters of a multivariate Gaussian distribution in a low-dimensional space (Methods and materials). Once the model has been trained, we view the location parameter of this distribution output by the encoder when fed a particular sequence as the embedding of the sequence in latent space. Viewing the embeddings of all the extant sequences in the alignment as a transformed version of the data, we introduce a model which assumes that they are generated by a multivariate Brownian motion along the phylogenetic tree within the low-dimensional latent space (Methods and materials). Note that this model is only introduced and applied to the embeddings obtained from the VAE after the latter has been trained in the standard fashion using stochastic gradient descent. The Brownian motion model applied to the embeddings of the extant sequences yields maximum likelihood estimates of “ancestral embeddings” at each internal node in the tree (Methods and materials). We then feed these estimated ancestral embeddings through the decoder of our trained VAE to obtain estimated ancestral sequences (Fig. 1, Methods and materials).

**Fig. 1:**
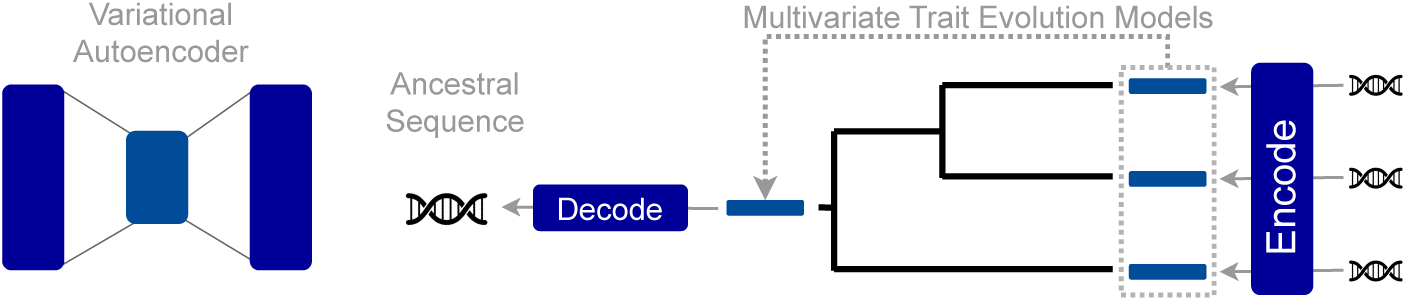
Graphical representation of method for ASR based on the embeddings from a VAE. The VAE is trained on extant sequences from a protein family to learn continuous representations of each extant member of the family. These “embeddings” are then used to fit a multivariate trait evolutionary model, yielding posterior estimates of “ancestral embeddings” for each internal node of the tree. Finally, the ancestral embeddings are fed back through the VAE decoder to arrive at reconstructions of the ancestral sequences.

We benchmark the VAE-based approach to ancestral sequence reconstruction (ASR) on sequences simulated to evolve in several ways. First, we consider the traditional model in which sites evolve independently in order to understand how VAE-based ASR does in this well-understood setting. We expected, however, that the advantage of an embeddings-based approach to ASR would only emerge in settings where the independent sites assumption is violated, as the embeddings would then be able to capture coevolutionary effects which standard phylogenetic models do not. To test this, we consider two epistatic models of sequence evolution: the Sequence Evolution with Epistatic Contributions (SEEC) model from (De La Paz et al. 2020), which amounts to Gibbs sampling from a Potts model, and the auto-regressive model of (De Leonardis et al. 2025), in which each site is simulated to evolve according to a continuous time Markov Chain whose stationary distribution depends on the sites that were evolved before it. Finding that a standard likelihood-based approach based on a model that assumes independent evolution of sites outperforms an embeddings-based approach even when applied to sequences simulated to evolve with epistasis, we are able to discount the utility of the embeddings-based approach for most, if not all, datasets. In the following sections, we describe the simulations results, first for the independent sites datasets, then the epistatic datasets.

## 4 Results

### 4.1 Independent site evolution

We first benchmarked the performance of VAE-based ASR when sequences evolve according to a standard model of protein evolution. We used the LG substitution model from Le and Gascuel (2008) to simulate multiple sequence alignments (MSA), where each site evolves independently of every other and considered three phylogenetic trees of various sizes on which to simulate evolution under this model. First, we considered tree topologies that were estimated to members of the Cluster of Orthologous Genes (COG) database (Tatusov 2001), taken from the benchmark set of FastTree (Price et al. 2009). In particular, we consider one tree with 1250 tips, inferred to the COG28 family, and one tree with 5000 tips, inferred to the COG2814 family, and on each of these two tree, we simulate independent-site evolution according to the LG model with Gamma distributed rate heterogeneity across sites (*α* = 0.5). Next, we generated a 10,000-tip tree with random uniform branch lengths between 0 and 0.3, and on this tree, we simulated evolution of sequences with the LG model and with equal substitution rates across sites, reproducing the tree and evolutionary simulation from Ding et al. (2019). In each case, the simulated sequences have 100 sites.

#### 4.1.1 Estimating ancestral embeddings in latent space

For each protein family, we trained variational autoencoders with 2- and 20- dimensional latent spaces on the collection of evolved sequences at the tips of the tree. We are able to directly visualize the 2-dimensional latent space, and we reproduce the star-shaped embedding manifold from Ding et al. (2019), as well as their finding that, at least in some cases, tracing the embeddings of a particular lineage of sequences in embedding space involves moving from the center of the latent space towards the periphery along the spikes in this star-shaped manifold (Fig. A1). The estimates of ancestral embeddings based on the Brownian motion model remain within this star-shaped manifold of embedding space (Fig. 2A) and, in particular, are relatively close to where the trained VAE embeds the corresponding ground truth ancestral sequence, which is known in our simulated setting (Fig. 2B). Averaging across ancestors, the relative distance ‖ *z − ẑ*‖*/* ‖ *z* ‖ between the embedding, *z*, of the ground truth ancestral sequence and its estimate based on the Brownian motion model, *ẑ*, was 0.18, 0.19, and 0.04 for the 2-dimensional VAEs and 0.30, 0.37, and 0.16 for the 20-dimensional VAEs, on the datasets with 1,250, 5,000, and 10,0000 sequences, respectively (Table A1).

**Fig. 2:**
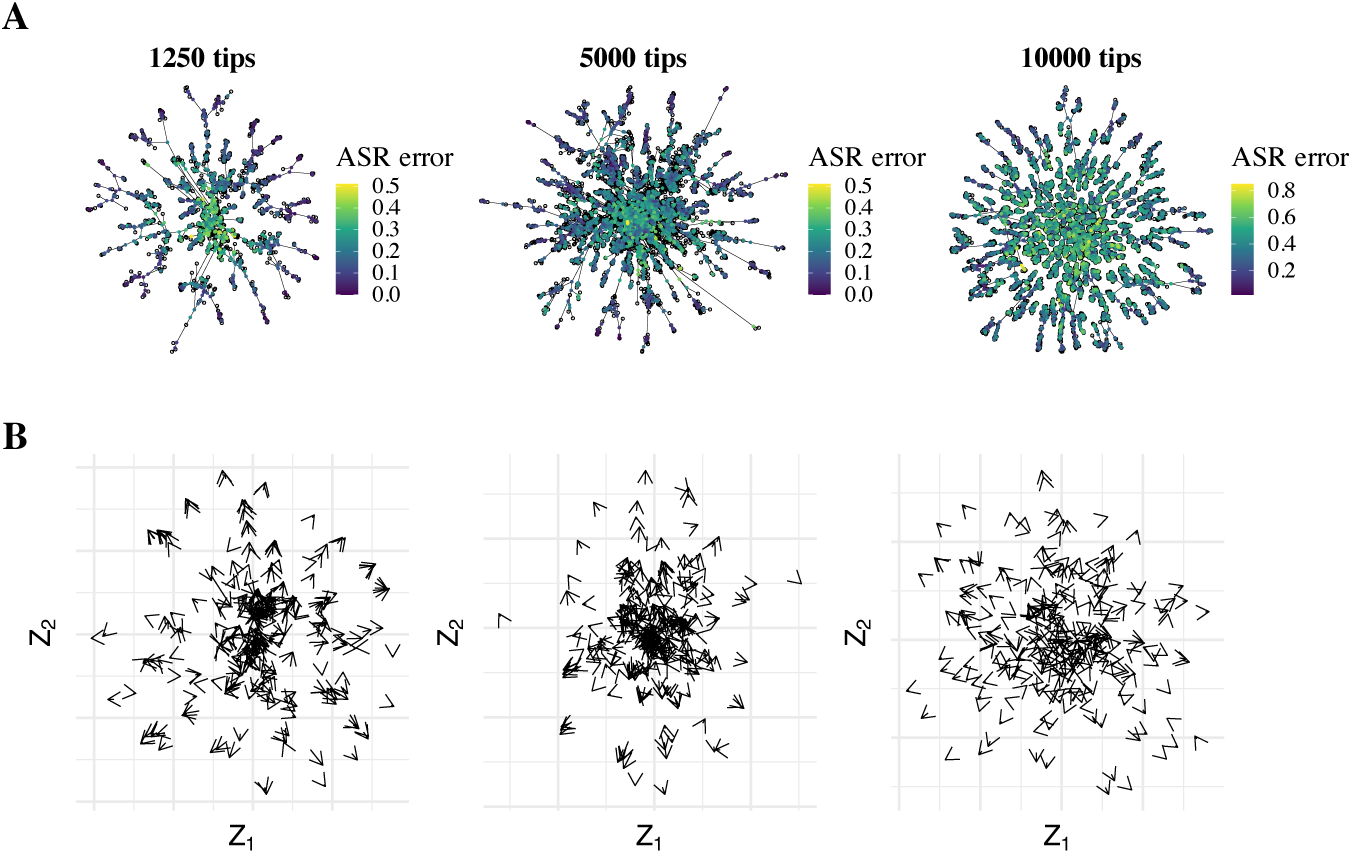
Visualizations of the latent space for independent sites simulations. Row **A**: The embeddings of all leaf sequences (unfilled dots) and the *estimated* ancestral embeddings based on the Brownian motion model (colorful, filled dots), with edge connections based on the phylogenetic tree. The colors of the estimated ancestral embeddings indicate the Hamming error of the VAE-based ancestral sequence reconstruction. In particular, the color of each estimated embedding represents the Hamming error between the ground truth ancestral sequence and the sequence obtained by feeding the estimated embedding through the VAE decoder. Row **B**: The error in estimating ancestral embeddings using the Brownian motion model. Each arrow starts at an estimated embeddings based on Brownian motion model and ends at the embedding of the actual sequence. To avoid over-plotting, a random subset of 300 ancestors are shown in each case. Column **1250 tips**: Family simulated with rate heterogeneity on COG28 tree. Column **5000 tips**: Family simulated with rate heterogeneity on COG2814 tree. Column **10000 tips**: Family simulated without rate heterogeneity on random, balanced tree from (Ding et al. 2019).

Even though interpolation with the Brownian motion model generally provided reasonable estimates of the embeddings of ancestral sequences, we observed a small but systematic bias on the periphery of the space, where ancestral sequences were often embedded further away from the origin than their descendant leaves. As a result, the estimates of ancestral embeddings based on phylogenetically interpolating the leaf embeddings were often shrunk towards the origin relative to the ground truth embeddings of the ancestral sequences, resulting in arrows in Figure 2B that point radially outwards at the periphery of the latent space. We quantified this by computing the average cosine of the angle between the ground truth embedding vector *z* and the error vector *z − ẑ*, finding that this average cosine value was positive for both the 2- and 20-dimensional VAEs (Table A1). While this artifact is certainly notable, we are ultimately interested in the reconstructed ancestral sequences obtained from decoding the estimated ancestral embeddings, not the estimated ancestral embeddings themselves. In section 4.2 we interrogate whether the error in the Brownian motion model’s estimation of ancestral embeddings is ultimately responsible for the error in the downstream ancestral sequence reconstruction.

#### 4.1.2 Benchmarking ancestral sequence reconstruction

In Figure 2A, the estimated ancestral embeddings are colored by the Hamming error that results from feeding them through the VAE’s decoder and comparing the resulting sequence to the ground truth ancestral sequence. We can see that the embeddings at the periphery of latent space are in fact decoded to more accurate estimates of the ancestral sequences than those in the center. Following the approach in De Leonardis et al. (2025), we additionally visualized ancestral sequence reconstruction accuracy for each trained VAE as a function of ancestral depth, which we define for our purposes as the patristic distance to the nearest leaf in the tree. In particular, given the set of all reconstructed ancestors in a particular family resulting from a particular method, we apply locally weighted scatterplot smoothing (LOWESS; Cleveland (1981)) to the scatter plot of ancestral reconstruction error versus ancestral depth (Methods and materials), and the purple curves in Figure 3 are the resulting smooths. Each panel of Figure 3 also includes horizontal lines at the average *self-reconstruction* error of the VAE on the collection of leaves on which it’s trained. In going from a 2-dimensional to a 20-dimensional latent space, the average self-reconstruction error made by the model on the sequences it’s trained on decreases, and this allows the method to achieve a lower ancestral sequence reconstruction error. Interestingly, for the larger datasets, the error in ancestral sequence reconstruction at every depth is lower than the VAE’s self-reconstruction error on the leaf sequences on which it is trained. We hypothesize that this is due to the tendency of VAEs to produce “blurry” reconstructions–that is, reconstructions that are biased towards an average of the model’s training distribution. Because an individual ancestral sequence resembles an average of leaf sequences, this regression to the mean in the VAE’s reconstructions actually causes, for these larger datasets, the ancestral sequences to be reconstructed with more accuracy than the self-reconstruction of the individual leaf sequences in the training distribution (see Appendix B for further discussion). For the 1250 sequence family, this is not the case, and looking at the 20-dimensional VAE’s performance on this family, we see that the self-reconstruction error on the leaf sequences, which is close to 0, resembles the error made in reconstructing the shallowest ancestors, suggesting that the model is learning to memorize the leaf sequences.

**Fig. 3:**
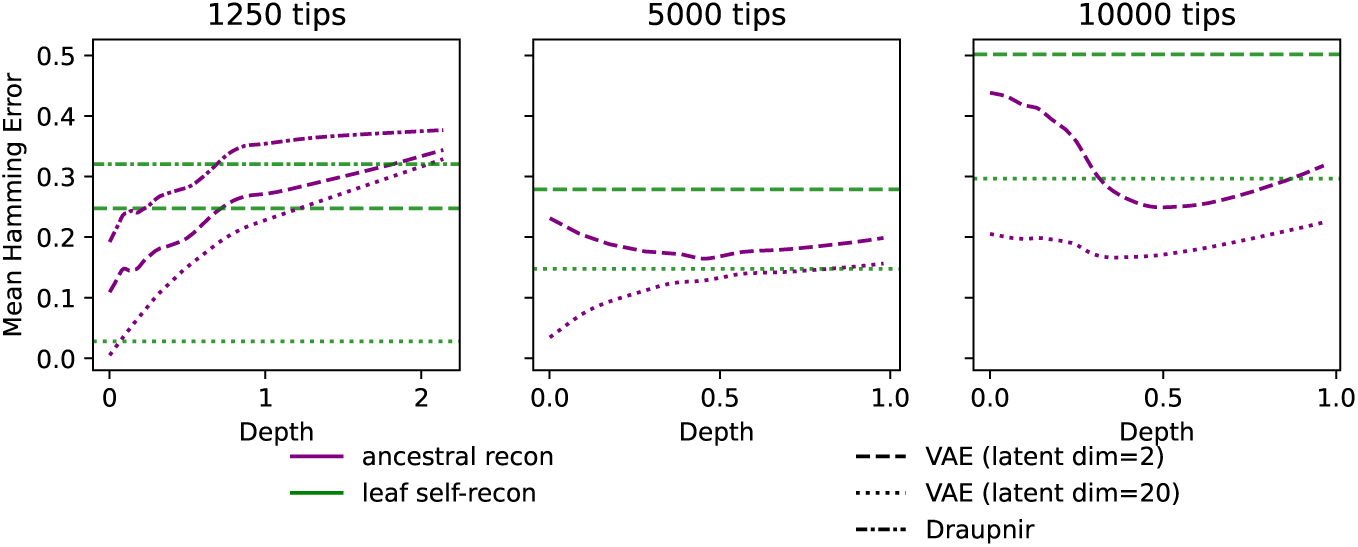
Sequence reconstruction error for different VAE-based approaches on three protein families simulated with independent evolution: **1250 tips**: family simulated with rate heterogeneity on COG28 tree; **5000 tips**: Family simulated with rate heterogeneity on COG2814 tree; **10000 tips**: family simulated without rate heterogeneity on random, balanced tree from Ding et al. (2019). Purple curves are LOWESS curves fit to the scatter plot of the method’s ancestral sequence reconstruction error versus the ancestor’s depth. Green horizontal lines are at the model’s error in self-reconstructing the leaf sequences, averaged over all leaves in the tree.

In addition to our VAEs, which do not incorporate a phylogenetic prior in the latent space, we also trained the “Draupnir” model from Moreta et al. (2022), which is a VAE that incorporates a phylogenetic prior, on the 1250 tip dataset. Draupnir’s inclusion of a phylogenetic prior on the embeddings when training the VAE does not result in improved self-reconstruction of leaf sequences, nor does it improve ancestral sequence reconstruction performance: our vanilla 2-dimensional VAE outperformed the Draupnir model at low depths and the 20-dimensional vanilla VAE outperformed the Draupnir model at every depth. Although the authors demonstrate the importance of the phylogenetic prior for inducing more phylogenetic structure in the organization of sequence embeddings in the latent space in a simulated dataset with 100 sequences, our results on this dataset with 1250 tips show that the phylogenetic prior does not improve the downstream ancestral sequence reconstruction. Indeed, Figure 2 shows that the embeddings organize according to phylogeny on our large datasets even without a phylogenetic prior based on the distribution of the sequences alone. Since the memory requirements of Draupnir scale quadratically with the number of sequences, we did not attempt running Draupnir on our larger MSAs with 5,000 and 10,000 leaves, but the above line of reasoning suggests that the phylogenetic prior would be even less necessary for these larger MSAs.

Moving to a comparison to non-VAE based methods, Figure 4 shows the ancestral reconstruction accuracy of the VAE-based approach using the model with a 20-dimensional latent space, along with the performance of several different existing methods: maximum likelihood estimation under a model of independent evolution using the IQTree software (Minh et al. 2020), maximum-likelihood estimation under the autoregressive model of evolution from De Leonardis et al. (2025), and maximum parsimony estimation (implementation details for each method in Methods and materials). As an absolute baseline, we also show the accuracy which results from using the consensus sequence from the alignment of leaf sequences (that is, taking the most frequent amino-acid at each position) as an estimate for each ancestor.

**Fig. 4:**
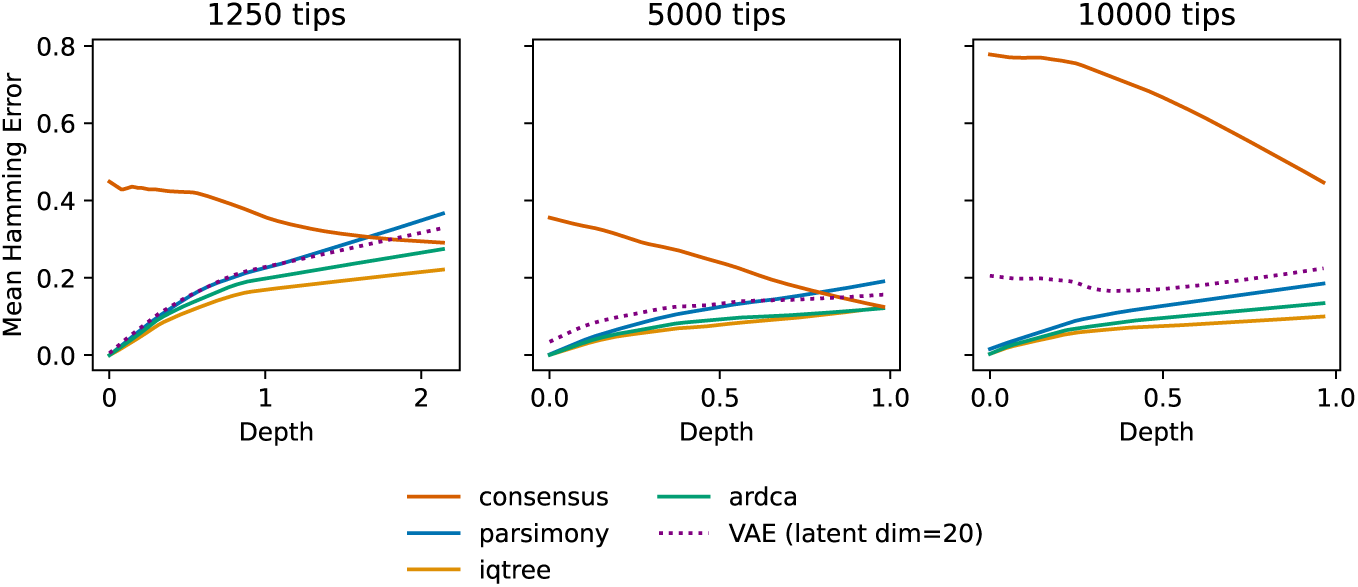
LOWESS curves fit to Hamming error versus depth-in-tree scatter plots for benchmark methods and for VAE-based approaches on each of the three protein families simulated with independent evolution. Benchmark methods include *consensus*: the consensus sequence from the MSA of the leaf sequences is used to estimate each ancestral sequence; *parsimony* : the maximum parsimony reconstructions of ancestral sequences; *iqtree*: marginal maximum likelihood reconstructions under an independent evolutionary model using IQTree; *ardca*: marginal maximum likelihood reconstructions based on the auto-regressive evolutionary model. **1250 tips**: family simulated with rate heterogeneity on COG28 tree; **5000 tips**: Family simulated with rate heterogeneity on COG2814 tree; **10000 tips**: family simulated without rate heterogeneity on random, balanced tree from Ding et al. (2019).

For each simulation setting, both likelihood-based methods for reconstruction, IQTree and ArDCA, dominate the VAE-based reconstructions at every evolutionary depth, and maximum parsimony dominates the VAE-based reconstructions for the 10,000 sequence family. Note that for these families simulated with independent sites, IQTree uses a correctly specified substitution model, and it is therefore unsurprising that it manages to achieve the best performance out of all the methods.

### 4.2 Epistatic evolution

To simulate epistatic evolution, we considered two distinct models, both based on transforming a generative model of the distribution of sequences in a protein family into a dynamic model of evolution. In both cases, we first fit the generative model to the Pfam family PF00072, a response regulator receiver domain. In the first case, we fit a Potts model to this MSA, and in the second case, we fit the auto-regressive model of Trinquier et al. (2021) (Methods and materials). To verify that epistasis is present in this family, we compute contact maps from each fitted model (Fig. A3, Methods and materials for implementation details).

To transform these generative models, Potts and auto-regressive, to models of sequence evolution, we follow the approaches of De La Paz et al. (2020) and De Leonardis et al. (2025), respectively. Specifically, for evolution with the Potts model, we performed Gibbs sampling from the Potts model along the phylogenetic tree, making the number of mutational steps on each branch proportional to its length. To simulate evolution with the auto-regressive model, we follow De Leonardis et al. (2025), who allow the evolution of each site in a sequence to depend on preceding ones (in some pre-specified order) by setting the equilibrium probabilities in the site’s Markovian dynamics at each branch in the tree to equal the predicted probabilities of the fitted autoregressive model given the previously evolved sites (see Methods and materials). In both cases, the tree on which we simulated evolution was the COG28 tree with 1250 leaves from the previous independent sites simulations.

Figure 5 displays the ancestral sequence reconstruction accuracy of VAEs with 2- and 20- dimensional latent spaces, as well as the other baseline methods, on the two epistatic families, as a function of ancestral depth in the tree. Although there is a switch, presumably due to the epistasis, in the best-performing method from IQTree to ArDCA, both of these methods continue to outperform the VAE-based approach across all depths, despite the fact that IQTree is maximizing the likelihood of a misspecified model with independent site evolution (the LG model, see Methods and materials). As mentioned, satisfactory performance from VAE-based ASR requires two distinct properties: the latent space must be phylogenetically structured, and the embeddings must preserve sufficient information about the sequences. Without the first property, it does not make sense to fit a phylogenetic Brownian motion model to the embeddings to estimate ancestral embeddings, and without the second, even if the phylogenetic Brownian motion model in the latent space is appropriate, feeding the interpolates representing ancestral embeddings back through the decoder will not result in accurate ancestral sequences. In our simulated setting where ancestral sequences are known, we can isolate the responsibility of the latter factor by simply considering the VAE’s ability to self-reconstruct ancestral sequences. We can then compare this inherent self-reconstruction inaccuracy to the overall ASR inaccuracy that results from first estimating ancestral embeddings with the Brownian motion model and then feeding these estimated ancestral embeddings through the VAE decoder.

**Fig. 5:**
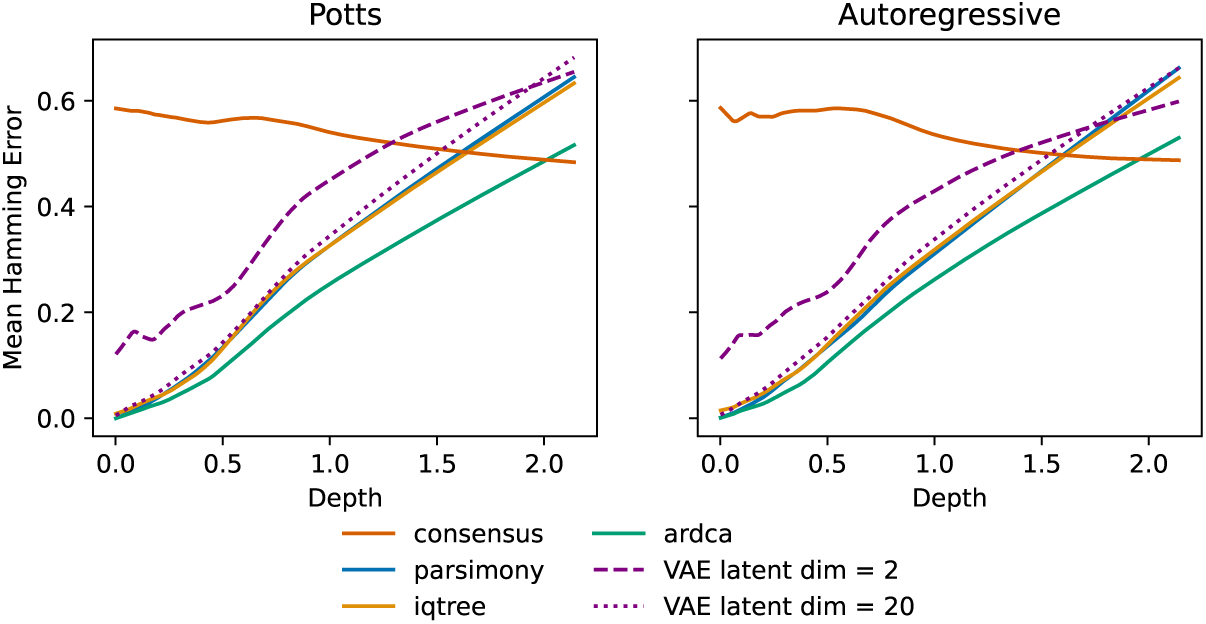
LOWESS curvers fit to Hamming error versus depth in tree for benchmark methods and VAE-based approaches on each of the epistaticly simulated families. See caption to Figure 4 for details on the benchmark methods included.

Figure 6 displays the result of this comparison. In particular, the red curves models the Hamming distance between ground truth ancestral sequences and their self-reconstructions, while the purple curves models the Hamming error from decoding the estimated embeddings, the same as the purple curves in Figure 5. In all cases, the Hamming error versus ancestral depth smooth for IQTree has been subtracted, so that the horizontal line at *y* = 0 represents exactly matching IQTree’s performance. Notably, the Hamming error from decoding the estimated embeddings tracks to a great degree the self-reconstruction Hamming error, with the red and purple curves crossing each other at various points but having the same general trajectories. For the VAE with a 20-dimensional latent space, the self-reconstruction of the ancestors is only slightly better than the accuracy based on decoding estimated ancestral embeddings for the Potts family, and is in fact slightly worse for the autoregressive family. In both cases, the self-reconstruction error sits above the line *y* = 0, meaning that IQTree is reconstructing ancestral sequences with higher accuracy than the VAE can self-reconstruct these ancestral sequences. What this suggests is that the ASR error made by the VAE-based approach is primarily driven by the inherent self-reconstruction error of the VAE on ancestral sequences, rather than error in the estimation of ancestral embeddings on the basis of the Brownian motion model.

**Fig. 6:**
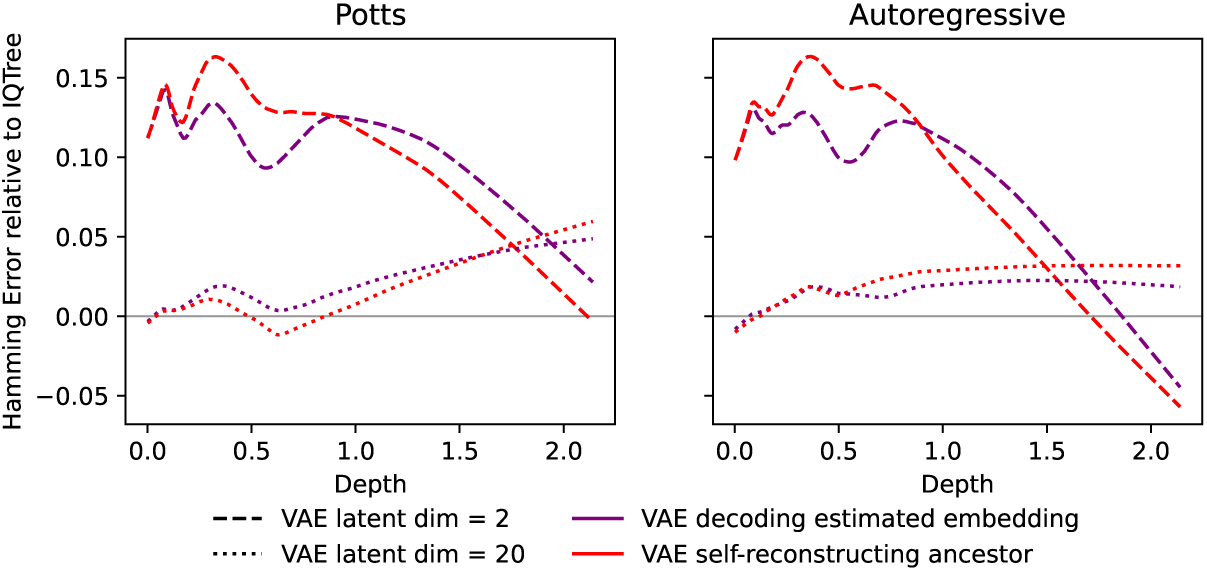
Comparison the error from decoding ground truth ancestral embeddings, i.e. self-reconstructing the ancestral sequences (red curves) versus decoding the ancestral embeddings *estimated* with the Brownian motion model (red curves). The LOWESS smooth of IQTree’s error from Figure 5 is subtracted from both, such that the line *y* = 0 represents matching the IQTree ASR error at every ancestral depth.

## 5 Discussion

In the VAE-based ancestral sequence reconstruction pipeline under study, we posit a Brownian motion model of evolution in the continuous latent space that is decoded by our learned VAE decoder to arrive at the evolution in sequence space. In this way, we avoid modeling evolution in the discrete sequence space which typically requires an assumed independence across sites to make the model computationally tractable. We hypothesized that we could more accurately capture the evolutionary traversal through sequence space in this way. In practice, however, we ran into difficulties. Our work highlights the importance of not uncritically assuming that deep-learning based embeddings can be used as the input data with which to perform common tasks in phylogenetics. Indeed, we show that interpolation in the latent space of a VAE trained in an unsupervised fashion on large protein families does not produce ancestral sequences that are as accurate as those reconstructed based on a substitution model’s likelihood, even when the substitution model is entirely misspecified, such as when sequences have evolved with epistasis.

We additionally show that the problem is not primarily that the embeddings from the VAE lack phylogenetic signal. In fact, Euclidean distances in the latent space do correlate with phylogenetic distances (Fig. A2), and the spikes in the latent manifold, as first observed in (Ding et al. 2019), do often correspond to clades in the phylogeny (Fig. A1). Rather, as we demonstrate in section 4.2, what is primarily responsible for the poor ancestral sequence reconstruction is simply the information loss inherent to the VAE: the VAE encoder loses too much information about the sequences in performing the compression to the latent space, and as a result, the embeddings cannot be decoded back to the original sequences with sufficient fidelity. This information bottleneck is less severe with a higher dimensional latent space, but the gains from increasing the latent dimension level off, and one must trade off the gains resulting from the higher dimensional latent space with an increased tendency towards overfitting to the leaf sequences and thus worse performance for deeper ancestors.

We stress that VAE embeddings can still be useful for visualizing, clustering, and exploring protein families despite falling short for ancestral sequence reconstruction, as we have documented here. More generally, embeddings from protein language model are extremely rich representations and have been demonstrated to be extremely effective for structural and functional prediction. One tantalizing if unrealized prospect is the incorporation of structural information into ancestral sequence reconstruction pipelines, and the mulitmodal protein language models which incorporate training on structural data provide an effective means to produce a shared embedding space for both sequence and structure (Fan et al. 2025; Heinzinger et al. 2024; Blaabjerg et al. 2024). Even with these exciting advances in developing more rich protein embeddings that results from both scaling up the training data and incorporating structural information, our study serves as a warning against uncritically employing embeddings as input data for tasks in which the goal is the fine-grained reconstruction of the evolutionary past.

## 6 Methods and materials

### 6.1 Trees

The random tree with uniform branch lengths between 0 and 0.3 was generated by calling the populate() method of the Tree class in python package ete3 (Huerta-Cepas et al. (2016), v3.1.1). The Cluster of Orthologous Genes trees from the FastTree study can be downloaded from http://www.microbesonline.org/fasttree/downloads/aa5Knew.tar.gz (trees with 5000 leaves) and http://www.microbesonline.org/fasttree/downloads/aa1250.tar.gz (trees with 1250 leaves). Price et al. (2009) describe how these unrooted trees were inferred to alignments of protein families from the Clusters of Orthologous Groups (COG), using either PhyML (for the 1250 leaf trees) or FastTree (for the 5000 leaf trees). Our simulations use specifically the COG28 tree (1250 leaves) and COG2814 tree (5000 leaves). Note that the FastTree algorithm results in some negative branch lengths in the 5000 leaf trees. Moreover, even the tree with 1250 leaves has several extremely short pendant edges, which would lead to duplicate evolved sequences. Therefore, we pre-processed the trees by (1) replacing all negative branch lengths with 0 and (2) subsequently trimming any pendant branch of length below 0.001. In order to simulate directed evolution along these trees, we rooted them at their midpoint.

### 6.2 Evolution simulations

#### 6.2.1 Independent sites

On the 10,000 sequence tree, we simulated evolution under the LG model without site rate heterogeneity to match one of the simulated MSAs from Ding et al. (2019). On the trees inferred to COG families, we use IQTree’s --alisim to simulate MSAs under the LG model with rate heterogeneity across sites. We set rates to vary according to a discrete (4 categories) Gamma distribution with shape parameter 0.5, -m LG+G{0.5}, representing sharp rate heterogeneity. For each of these families, sequences had 100 sites.

#### 6.2.2 Potts

Our first simulation of epistatic evolution was performed by Gibbs sampling from a Potts model along a phylogeny. Evolution proceeds by randomly choosing a site in the sequence to mutate and sampling from the conditional distribution of this site given all the other sites according to the Potts model (note that this may result in no change if the sampled amino acid is the same as the current one). The number of times we do this on a given branch equals the branch length times the length of the sequence. We rely on the implementation in the Julia package PottsEvolver.jl (Pagnani and Barrat-Charlaix 2025). The Potts model under which evolution is simulated was trained with a Boltzmann machine to reproduce the statistics of an MSA of transporter receiver domains (PF00072) using the Julia implementation of adabmDCA 2.0 (Rosset et al. 2025). To compute a contact map from this trained model, we use this package to compute the average-product corrected Frobenius norms of the coupling matrices.

#### 6.2.3 Autoregressive

The autoregressive model of evolution is formulated in De Leonardis et al. (2025). We fit an autoregressive generative model to PF00072 using the ardca function in the Julia package ArDCA.jl (Trinquier et al. 2021). To compute the contact map from this fitted model, we use the method epistatic score from this package, which computes effective couplings from the autoregressive model and then computes the average-product corrected Frobenius norm of the coupling matrices. To simulate evolution using the parameters from the fitted model, we rely on the function ASR.Simulate.evolve from AncestralSequenceReconstruction.jl (De Leonardis et al. 2025).

### 6.3 Real protein families

#### 6.3.1 Response regulator receiver domain (PF00072)

We take the alignment for PF00072 constructed in Jones et al. (2011), which includes 2 74,836 sequences with 126 sites. In order to remove excessively gappy sequences and positions, we performed the following steps in order:

1. We remove any sequences that were more than 20% gaps. 7
2. We remove any positions that are at this point more than 20% gaps. 9

This resulted in an MSA with 72,458 sequences with 114 sites, to which we fit the 1 generative models, Potts and autoregressive.

#### 6.3.2 Beta lactamase (PF00144)

We downloaded the MSA for PFAM family PF00144 from the Github repository released with Detlefsen et al. (2022), accessible at https://github.com/MachineLearningLifeScience/meaningful-protein-representations/blob/master/tape/PF00144full.txt. We then applied the same filtering steps mentioned above, reducing the length of the MSA from 2592 to 229 and the number of sequences from 36,328 to 23,698.

A phylogenetic tree was inferred from this pre-processed MSAs using IQTree2 (Minh et al. 2020). Because no prior information was available regarding the appropriate evolutionary model, we enabled automatic model selection using the -m MFP option, which evaluates a range of candidate substitution models and selects the best-fitting model based on the lowest Bayesian Information Criterion (BIC).

The discussed subset of 134 representative sequences are the intersection of the 200 sequences used for ancestral reconstruction in Detlefsen et al. (2022) and the sequences that were retained in our pre-processed MSA.

### 6.4 Variational Autoencoder

A variational autoencoders (VAE) is a neural network with an autoencoder architecture that is trained to learn a probabilistic, latent variable model of the data it encodes (Kingma and Welling 2013). Autoencoder neural networks generally consist of two subnetworks: an encoder and a decoder. The encoder network learns a mapping from the high-dimensional space of the data, in our case, protein sequences, to a lower-dimensional embedding space. The decoder, in turn, learns to map the lower dimensional space back to the original space in such a way that it approximates the inverse of the encoder. Autoencoders thus learn lower-dimensional representations of the data they are trained on. In a variational autoencoder, in order to avoid overfitting and produce a smoother embedding space, a prior is placed on the embeddings and the autoencoder is trained to learn a probabilistic model of the data.

Formally, for each sequence *s* in our alignment, it is assumed that a continuous, low-dimensional latent variable *z* first gets generated according to a simple distribution *p*(*z*); then *z* gets mapped in a complex fashion to a distribution *p**_θ_***(*· | z*) over sequences, where the mapping is parameterized by unknowns ***θ***; finally, *s* is sampled from this distribution. The decoder, DEC, of the autoencoder is used to execute this complicated mapping from *z* to *p**_θ_***(*· | z*). If we were to attempt to learn the weights and biases, ***θ***, of the decoder by directly maximizing the marginal likelihood of the data, we would encounters the intractable integrals in

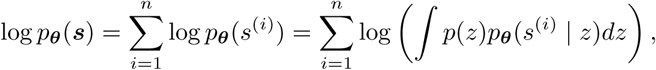

where *n* is the total number of sequences in the alignment ***s*** = {*s*^(1)^*, s*^(2)^*, …, s*^(*n*)^}. To facilitate the learning of this model, we introduce the encoder neural network,

ENC, parameterized by ***ϕ***, which should map each sequence *s* to a distribution that approximates the posterior of the latent variable given the sequence, ENC(*s*) = *q**_ϕ_***(*· | s*) *≈ p****_θ_***(*· | s*). To simultaneously train the encoder and decoder, we consider, instead of log *p****_θ_***(***s***), the evidence lower bound (ELBO) objective function

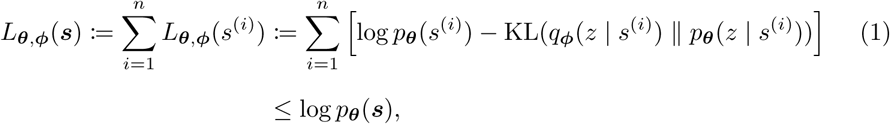

where KL(*·* ‖ *·*) denotes the Kullback-Leibler (KL) divergence and the inequality is due to the non-negativity of the KL divergence.

Maximizing (1) jointly with respect to both ***θ***, the parameters of the decoder, and ***ϕ***, the parameters of the encoder, simultaneously targets large log *p**_θ_***(***s***), which corresponds to the original goal of finding a good generative model, and small 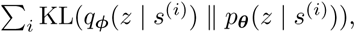, which corresponds to accurate inference of the latent variables for the sequences, given our generative model. However, if the only way to make the KL(*q**_ϕ_***(*z | s*^(*i*)^) ‖ *p****_θ_***(*z | s*^(*i*)^)) small involves choosing a ***θ*** for which log *p****_θ_***(***s***) is small, there can be a tradeoff between the two competing objectives. In order to avoid this situation, we use a sufficiently flexible neural network for the encoder, so that *s* ↦ *q****_ϕ_***(*· | s*) is flexible enough to approximately match *s* ↦ *p****_θ_****∗* (*· | s*), where ***θ****^∗^* = arg max***_θ_*** log *p****_θ_***(***s***).

Unlike the marginal log-likelihood, the ELBO objective can be tractably estimated, since we have

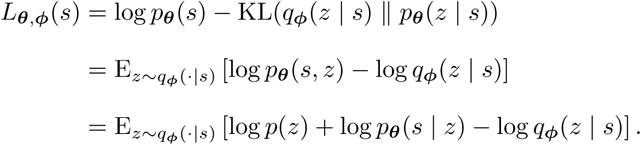

We can calculate the expression appearing inside the expected value for any *z* and thereby obtain a stable Monte Carlo estimate of our new objective evaluated at a single training sequence *s* by sampling *z* from *q**_ϕ_***(*· | s*) and plugging into the expression. Estimates of the gradients of the negative ELBO with respect to ***θ*** and ***ϕ*** can then be obtained with autodifferentiation in tandem with the reparameterization trick, and we can use these estimated gradients to optimize *L****_θ,ϕ_***(***s***) with mini-batch stochastic gradient descent.

In our implementation of a vanilla VAE for protein sequences of fixed-length *m*, the likelihood is a product of independent categorical distributions over amino acids for each position in the sequence,

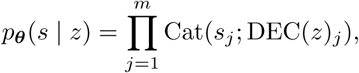

where DEC(*z*)*_j_* is the probability distribution over amino acids at the *j*th position in the sequence computed by the decoder. Meanwhile, is a standard normal *p*(*z*) = *N* (*z*; **0***_d_,* ***I****_d_*), where *d* is the dimension of the latent space, and the variational approximation to the posterior of the latent variable is

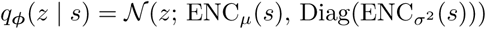

Finally, following Ding et al. (2019), our encoder and decoders are both given by multi-layer perceptrons, each with a single hidden layer:

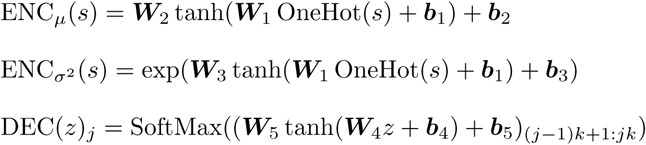

for *j* = 1, 2 *…, m*. Here, *k* is the number of different amino acids and equals 21 and 20, for the alignments with and without gaps, respectively. We use a hidden dimension of 500 (1000) in models fit to simulated (real) MSAs and use a latent dimension of 2 and 20 to explore the latter hyperparameter’s impact on the results. In line with the clustering results from Ding et al. (2019), we observed diminishing returns to increases in the latent dimension, and the improvement plateaus by the time the latent dimension is 20.

When initially training the VAEs, we held out 10% of the extant sequences from the MSA to use as a validation set. We used weight decay to regularize our networks, and all of the models shown in the paper were fit with the weight decay parameter that achieved optimal importance weighted ELBO (IW ELBO, Burda et al. (2015)) on the held out set (Table 1), which plateaued for all models by 500 epochs of training. Upon finding this suitable weight decay value, we fit the VAE with the full set of extant sequences, and used this latter model, trained on the full set of leaf sequences, to perform the ancestral sequence reconstruction.

**Table 1:** VAE performance across all datasets, measured by Importance Weighted (IW) ELBO per site in a held-out set of leaf sequences. The IW ELBO is computed with 5000 samples. Note that the non-independent evolution simulators and real MSA included gap characters, but the independent model (LG) did not, so the IW ELBO values are not directly comparable.

| Tree | Evolution simulation | VAE latent dimension | Weight decay | IW ELBO/<br>Site |
| --- | --- | --- | --- | --- |
| COG28<br>(1250 tips) | Independent | 2 | 5e-3 | -1.15 |
|  |  | 20 | 1e-2 | -1.08 |
|  | Potts | 2 | 5e-3 | -1.36 |
|  |  | 20 | 1e-2 | -1.28 |
|  | Autoregressive | 2 | 5e-3 | -1.40 |
|  |  | 20 | 1e-2 | -1.33 |
| COG2814<br>(5000 tips) | Independent | 2 | 1e-3 | -1.15 |
|  |  | 20 | 5e-3 | -1.13 |
| Random tree<br>(10000 tips) | Independent | 2 | 1e-3 | -1.92 |
|  |  | 20 | 5e-3 | -1.80 |
| Inferred<br>(23698 tips) | Real MSA<br>(PF00144) | 2 | 0 | -1.55 |
|  |  | 20 | 5e-3 | -1.44 |

We explored adding additional hidden layers to the encoder and decoder, but found that these changes did not result in improvements in the IW ELBO on the held out set, nor to improvements in performance on the downstream task of ASR.

### 6.5 Maximum likelihood ancestral reconstruction under Brownian motion

Once the VAE is trained, embeddings for each of the tips of the phylogenetic tree are obtained by passing each extant sequence through the encoder ENC*_µ_*, resulting in a matrix

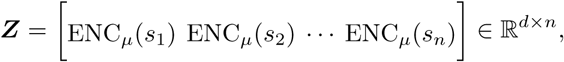

where *n* is the number of tips in the tree and *d* is the latent dimension of the VAE. For each internal node *i* in the tree, an “ancestral embedding” is estimated under an assumed model of Brownian motion as follows. Let *C_i_* be the phylogenetic covariance matrix under Brownian motion that would apply to the tips given a tree rooted at *i*, i.e. assuming trait evolution begins at *i* and is directed away from *i*. Formally, [*C_i_*]*_ab_* gives the distance from *i* to the most recent common ancestor of tips *a* and *b* in the tree rooted at *i*. Then the maximum likelihood estimate of the ancestral embedding *z_i_* is given by

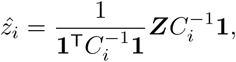

where **1** *∈* R*^n^* is a column vector of ones. The quadratic forms with *C^−^*^1^ appearing in this equation can be computed efficiently (linearly in the number of taxa) with the algorithm from Tung Ho and Ané (2014). We rely on the implementation in the R package RPhylopars (Goolsby et al. 2016).

### 6.6 Ancestral Sequence Reconstruction with the VAE

To reconstruct the sequence sequence for ancestor *i*, we feed its estimated ancestral embedding *ẑ_i_* through our trained VAE’s decoder and then, at each position, select the amino acid assigned the highest probability:

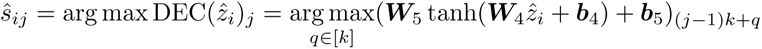

for *j* = 1, 2 …,*m*.

### 6.7 ASR Benchmark Methods

#### 6.7.1 Draupnir

We trained the Draupnir VAE on the MSA of 1250 sequences simulated with independent sites evolution on the COG28 tree. We tried training with both MAP (no encoder, latent variables are themselves optimized to maximize the joint distribution) and full variational inference, which resulted in similar results, and we only show the results from MAP inference. For each ancestor, reconstructed sequences are obtained by taking the most likely amino acid for the ancestor at each position.

#### 6.7.2 Maximum parsimony

We apply Fitch’s algorithm (Fitch 1971) to recover the assignment of amino acids to ancestral nodes that minimize the total number of changes along the tree. If there are multiple possible choices that achieve this minimal changes, we choose one arbitrarily to evaluate for Figures 4 and 5. For the reconstruction of the root of the beta lactamase family in Figure C1, we assign equal probabilities to each such solution.

#### 6.7.3 IQTree

We reconstruct likelihood-based estimates of ancestral sequences by running IQTree with the -asr flag. We impose the LG model with rate heterogeneity (-LG+G) even for the datasets that were simulated with epistatic evolution. We fix the tree topology and do not have IQTree optimize the branch lengths (-blfix). IQTree outputs the posterior distribution over amino acids at each site in each ancestral node in the tree, and we choose for each position the amino acid assigned the greatest probability to obtain IQTree’s reconstructed ancestral sequence. Note that IQTree does not include gap characters in its vocabulary, so no posterior probability is assigned to gaps.

#### 6.7.4 Autoregressive reconstruction

To infer ancestral sequences under the autoregressive model from Bisardi et al. (2021), we rely on the algorithm from De Leonardis et al. (2025) implemented in AncestralSequenceReconstruction.jl. We fit the autoregressive model that is required for the algorithm to the alignment of leaf sequences. We keep the tree topology and its branch lengths fixed when reconstructing ancestors with the infer_ancestral() function in AncestralSequenceReconstruction.jl. For Figures 4 to 6, we choose the maximum a posteriori estimate to evaluate, but AncestralSequenceReconstruction.jl can also provide samples from the posterior, and we display the distribution of these posterior samples for the reconstruction of the root of the beta lactamase family in Figure C1.

### 6.8 LOWESS curves

For the set of reconstructed ancestral sequences from each method, a LOWESS curve was fit to the scatter plot of Hamming error versus ancestral depth (defined as patristic distance to nearest leaf) using the function lowess from the module statsmodels.nonparametric.smoothers_lowess in Python. We set frac, the fraction of the data used for each local fit, to be 0.3.

## 7 Data Availability

The raw and pre-processed multiple sequence alignments for PF00072 and PF00144 are made available at the Github repository for this paper: https://github.com/solislemuslab/asr-ae/. This repository also contains the code for the simulation of MSAs under different models of evolution, the training of the VAEs, and the benchmarking of the embeddings-based approach to ASR against other standard methods.

## 8 Acknowledgements

This work was supported by the National Science Foundation (DEB-2144367 to CSL). This material is based upon work support by the National Institute of Food and Agriculture, United States Department of Agriculture, Hatch project 7007384 (to CSL).

CSL participated in the Institute for Computational and Experimental Research in Mathematics in Providence, RI “Theory, Methods, and Applications of Quantitative Phylogenomics” program and presented early versions of this work.

## Appendix A Additional Tables and Figures

**Fig. A1:**
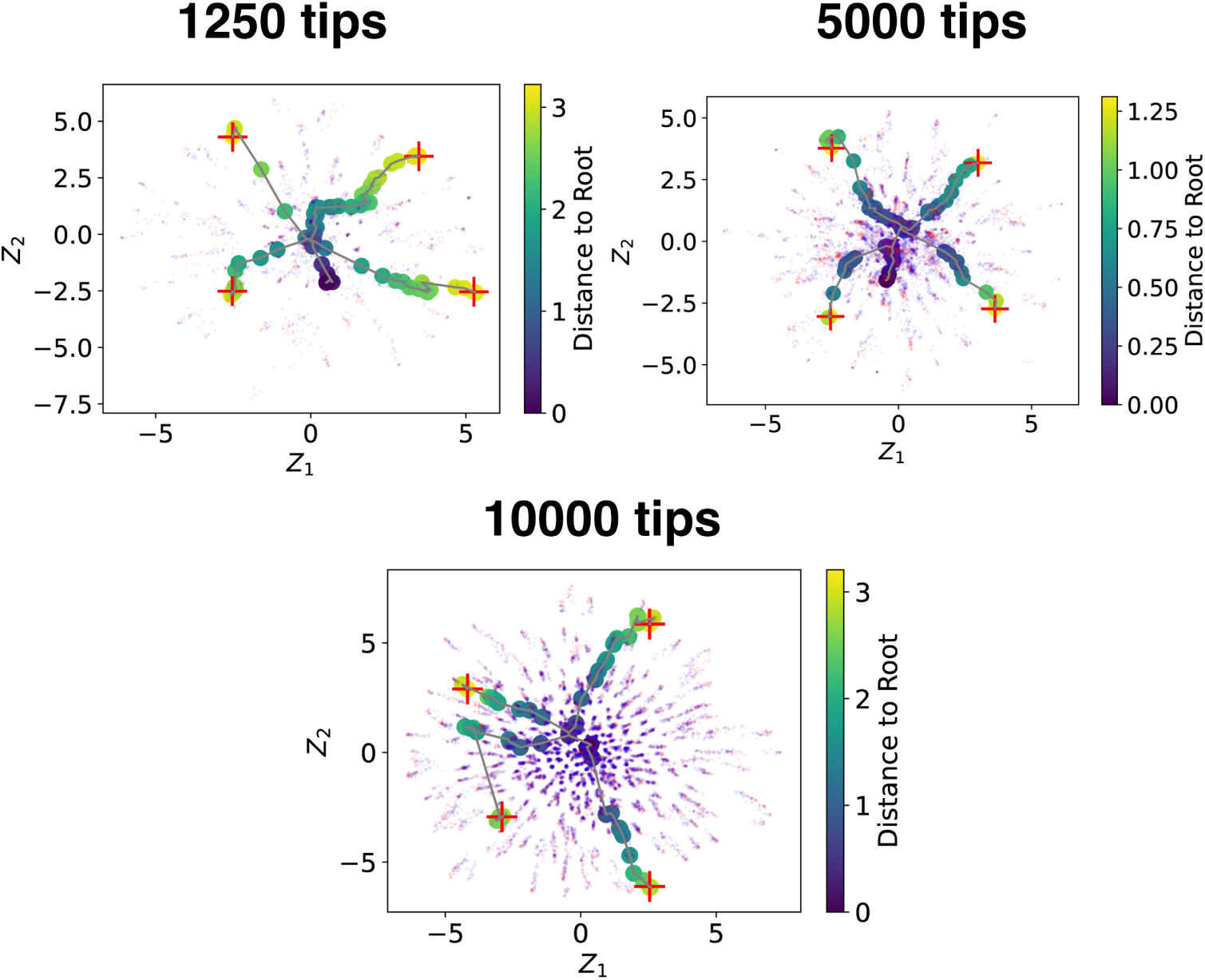
Embeddings of all leaf (blue dots) and *ground-truth* ancestral (tiny red dots) sequences from the trained VAE. The embeddings of four leaves at the outskirts of the embedding space are singled out with red crosses. The embeddings of the ancestral lineage of each singled-out leaf is connected by a line, and these ancestral lineages coalesce at the root in the center of embedding space, as in Ding et al. (2019). The colors of the embeddings of the ancestors of the singled-out leaves are based on their distance to the root.

**Table A1:**
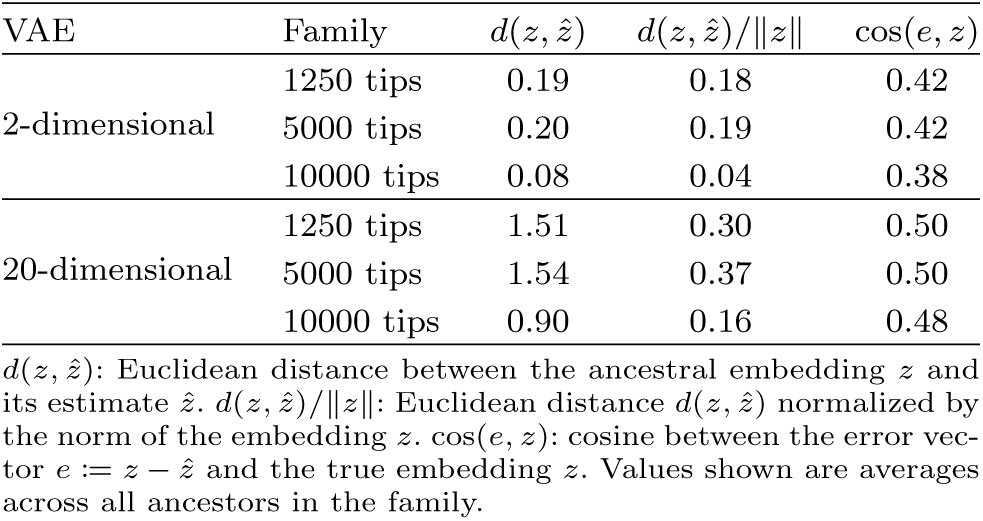
Summary statistics on ancestral embedding estimation for simulations with independent sites.

**Fig. A2:**
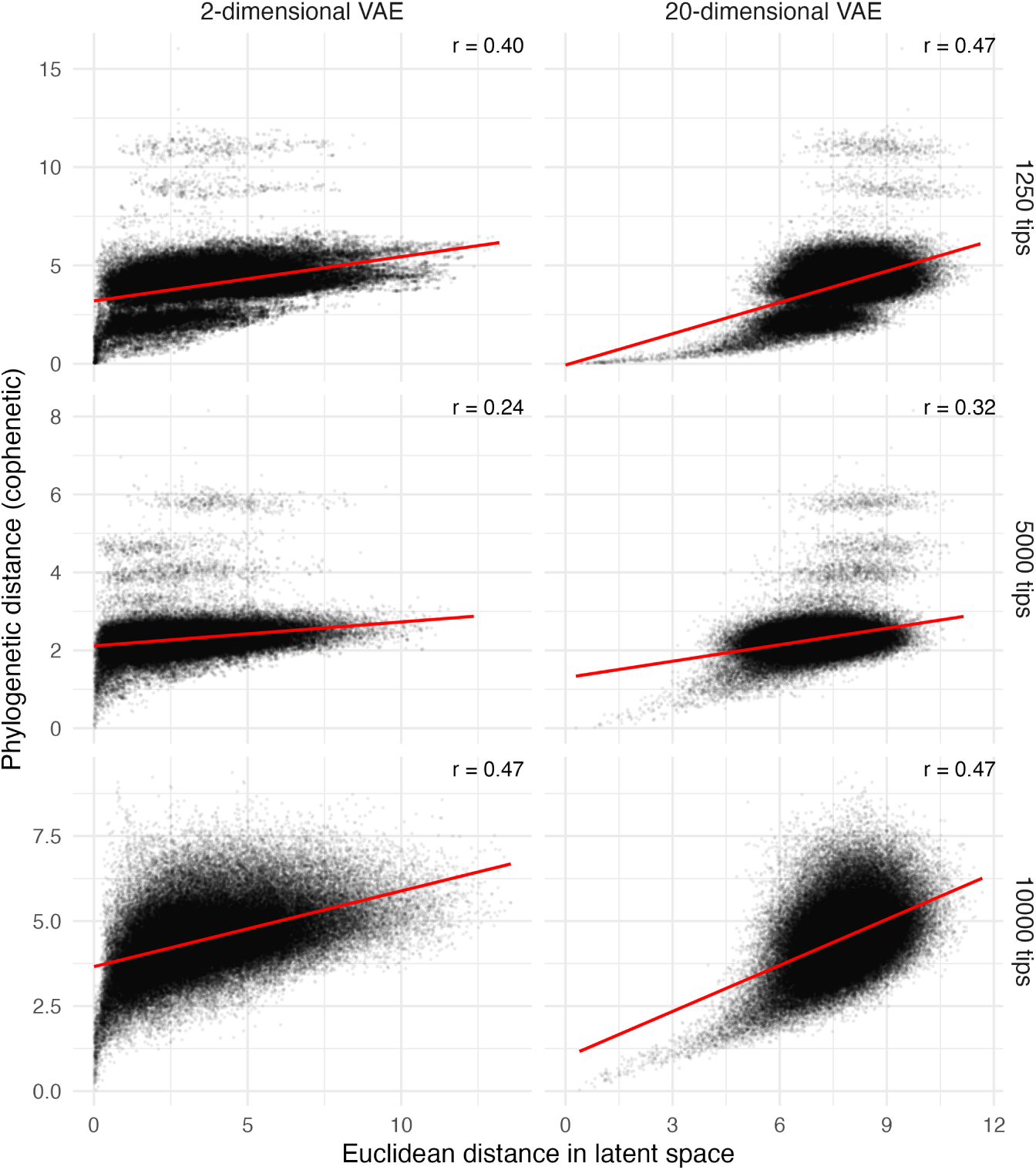
Scatter plots of the phylogenetic_39_distance (in branch lengths) between pairs of leaves versus the Euclidean distance between their embeddings for the independently simulated MSAs. In each case, 500 leaves are randomly sampled and pairwise distances are shown for these 500 leaves.

**Fig. A3:**
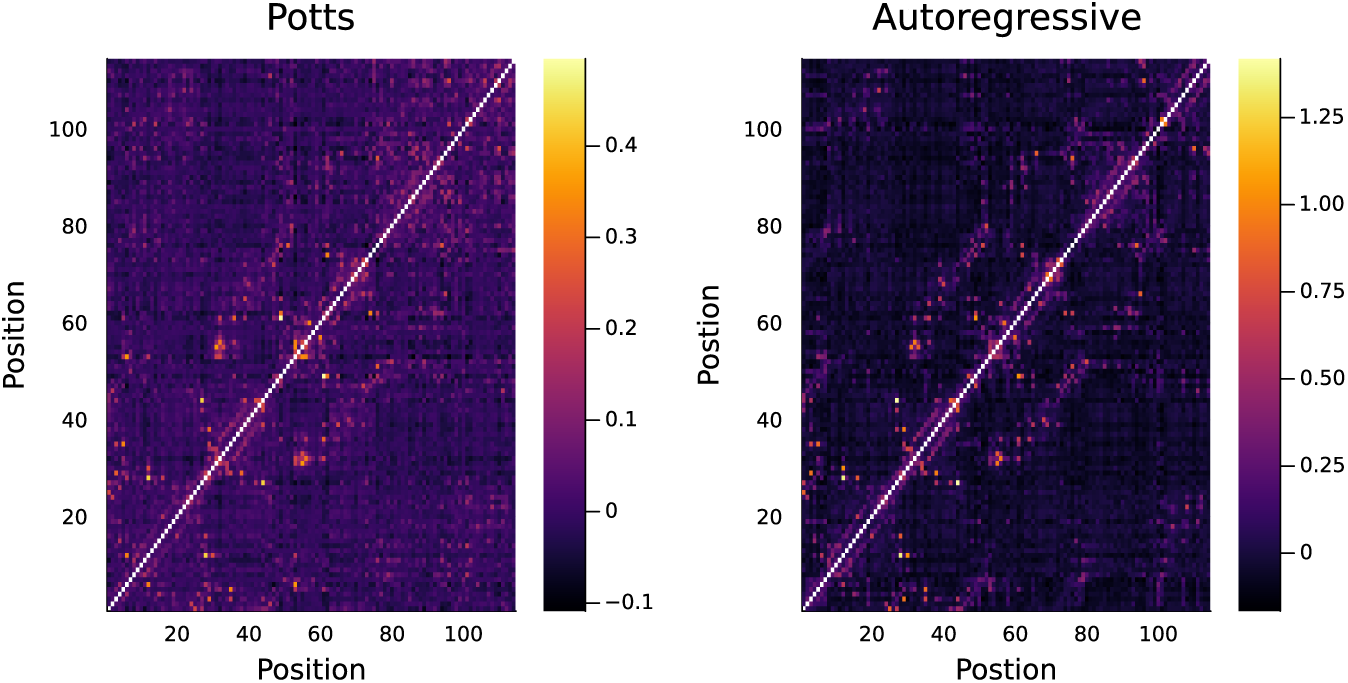
Contact maps from different generative models fit to multiple sequence alignment of response regulator receiver domain (Pfam family PF00072) Left: Potts model; right: autoregressive model.

**Fig. A4:**
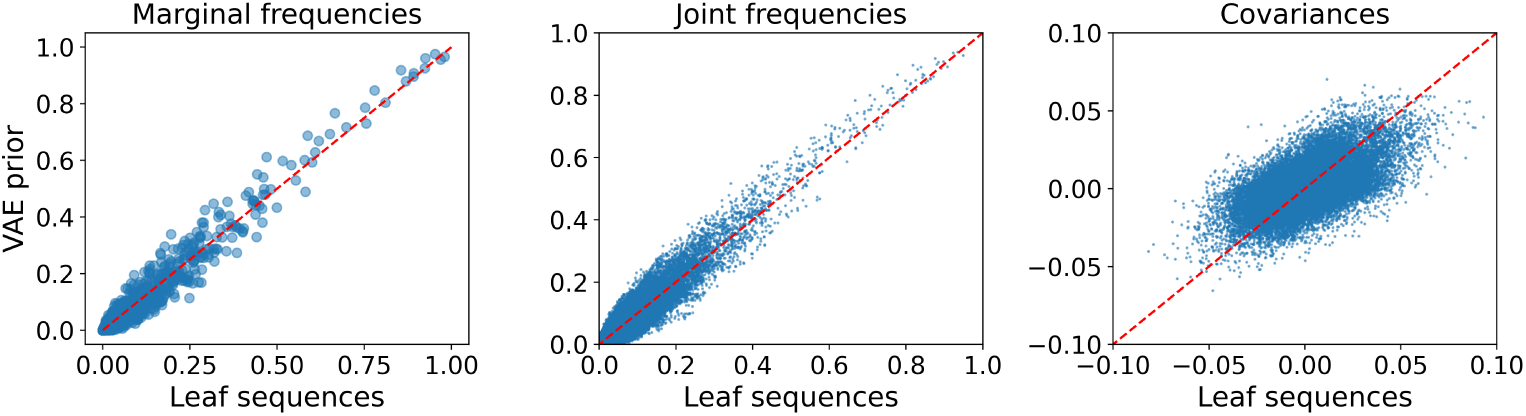
Statistics calculated on sequences sampled from the VAE with a 2-dimensional latent space fit to the MSA simulated by sampling from the Potts model along the COG28 tree, versus the corresponding statistics calculated from the MSA itself. Covariances are joint frequencies minus the product of corresponding marginal frequencies.

## Appendix B Leaf sequence self-reconstruction error

There is variability in the model’s ability to self-reconstruct different leaf sequences which is hidden by the horizontal lines at the average self-reconstruction error across all the leaves in the tree in Figure 3. In particular, some leaf sequences are distant from the bulk of the distribution which the VAE learns to model, and the model performs worse at self-reconstructing these. Figure B1 shows ancestral sequence reconstruction error and leaf self-reconstruction error both as a function of depth in the same plot, where, for the purposes of this plot, we define the “depth” of a leaf as its distance to the nearest leaf in the tree that is not itself. In our 1250-sequence and 5000-sequence families, the maximum leaf depth is actually much larger than the maximum depth of an ancestor, resulting from a few extremely long pendant edges in their trees. We can see that, for all but the 20-dimensional VAE applied to the 1250-sequence family, the error in the self-reconstruction of leaves in fact exceeds the ancestral reconstruction error not just on average, but at each particular depth, with the two converging as depth approaches 0.

**Fig. B1:**
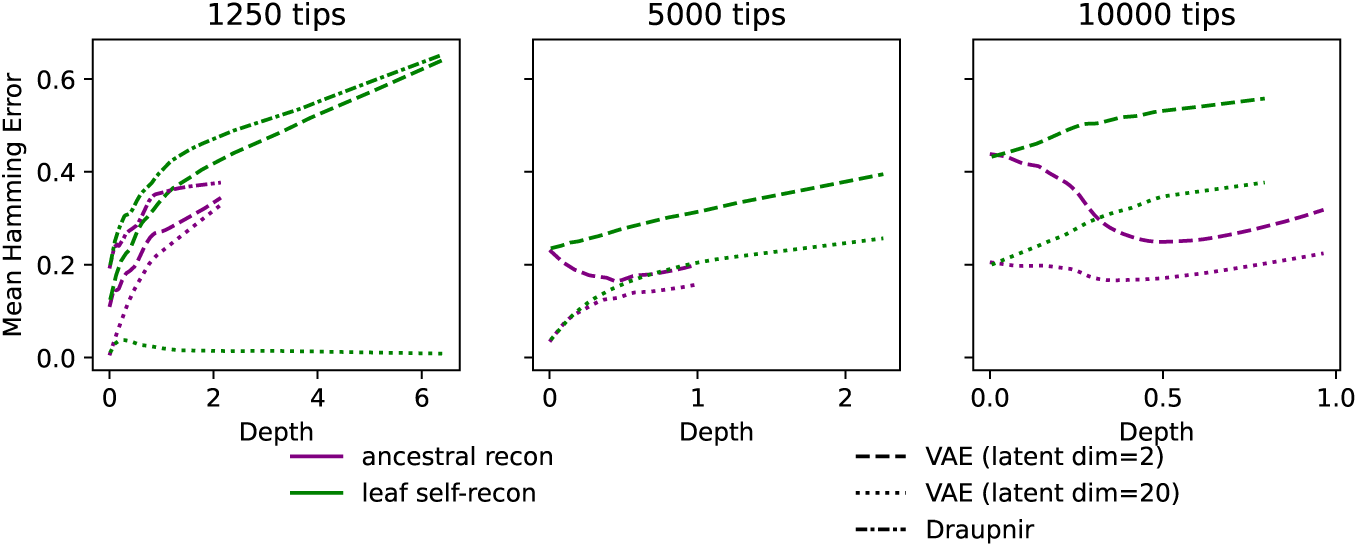
Sequence reconstruction error for different VAE-based approaches on three protein families simulated with independent evolution: **1250 tips**: family simulated with rate heterogeneity on COG28 tree; **5000 tips**: Family simulated with rate heterogeneity on COG2814 tree; **10000 tips**: family simulated without rate heterogeneity on random, balanced tree from Ding et al. (2019). Purple curves are LOWESS curves fit to the scatter plot of the method’s ancestral sequence reconstruction error versus the ancestor’s depth. Green curves are LOWESS curves fit to the scatter plot of the method’s sequence self-reconstruction error on leaves versus the leave’s depth, defined as distance to nearest leaf in the tree other than itself.

## Appendix C Beta-lactamase

We additionally ran our pipeline on the Beta-lactamase family (PFAM family PF00144, Mistry et al. (2020)). We trained the VAE model with latent dimensions of 2 and 20 on a pre-processed MSA that had 23,698 sequence with 229 sites (see Methods and materials). On the basis of this same MSA with 23,698 tips, we inferred a tree using IQTree. We then took the subtree induced by a subset of 134 representative sequences from Detlefsen et al. (2022) (see Methods and materials), computed their embeddings by applying the trained VAE encoder, reconstructed ancestral embeddings for the internal nodes of the subtree using Brownian motion, and decoded these estimated ancestral embeddings to obtain estimated ancestral sequences. Figure C1 shows the predicted distribution of amino acids in the first 20 positions of the ancestral sequence at the root according to different methods, including both VAE models, as a probability logo plot. Embeddings from the 2 dimensional VAE and reconstruction of a different ancestral sequence are shown in Figures C2 and C3, respectively. Overall, we see that the uncertainty in the VAE-based reconstructions tracks the uncertainty from other methods; however, there are discrepancies between the VAE-based reconstructions and the other methods. For example, in position 6, parsimony, IQTree, and ArDCA are unanimous in reconstructing a glutamic acid, while both VAEs are quite uncertain with the low-dimensional VAE assigning the highest probability to glutamine and the high-dimensional VAE assigning the highest probability to alanine.

**Fig. C1:**
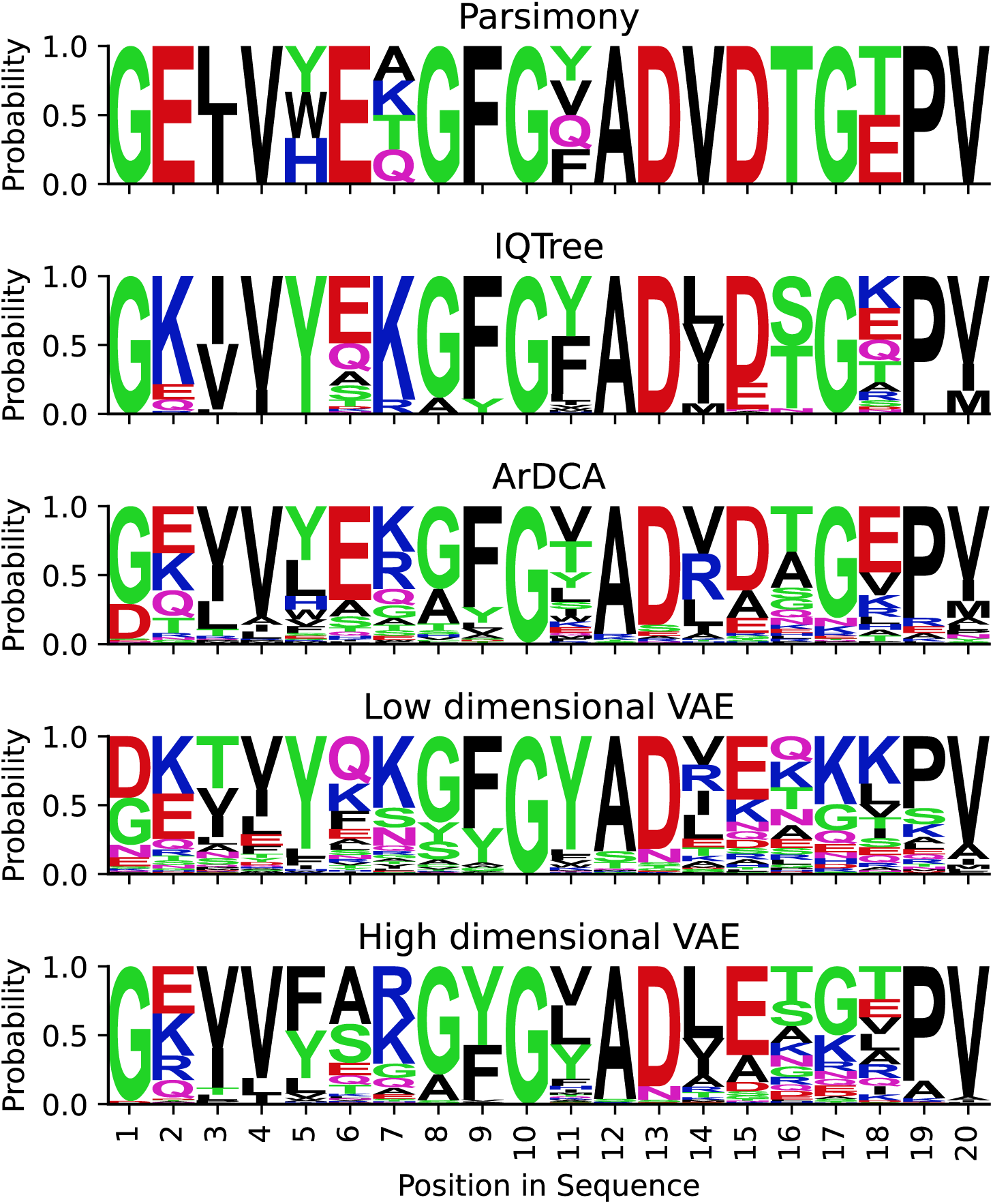
Probability logo plot of reconstructions of first 20 positions of root sequence of the Beta-lactamase family (PF00144) with different methods: maximum parsimony (Parsimony), maximum likelihood under an independent sites model (IQTree), maximum likelihood under the autoregressive evolutionary model (ArDCA), a VAE with a 2-dimensional latent space (Low dimensional VAE), and a VAE with a 20-dimensional latent space (High dimensional VAE).

**Fig. C2:**
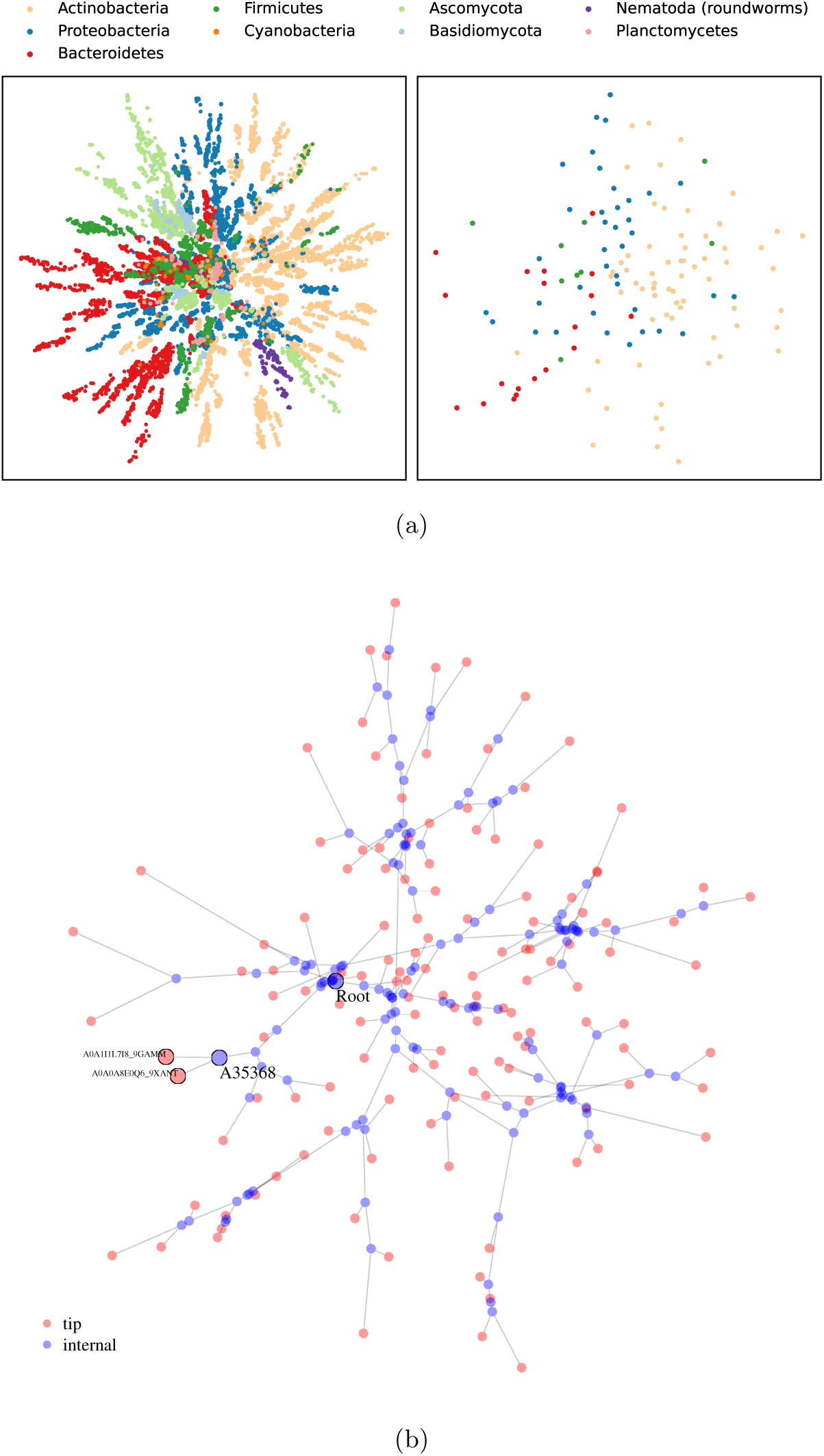
(a) Embeddings of sequences from the PF00144 MSA colored by phylum of species that sequence is sampled from. Left: embeddings of all sequences. Right: embeddings of representative sequences from Detlefsen et al. (2022) (b) Embedding of inferred phylogenetic tree in latent space. Positions of internal nodes in blue are estimated under assumed Brownian motio^4^n^4^ along the phylogeny. Root is highlighted, as well as a shallow ancestor, A35368, and its two leaf children, A0A1I1L7I8 9GAMM and A0A0A8E0Q6 9XANT.

**Fig. C3:**
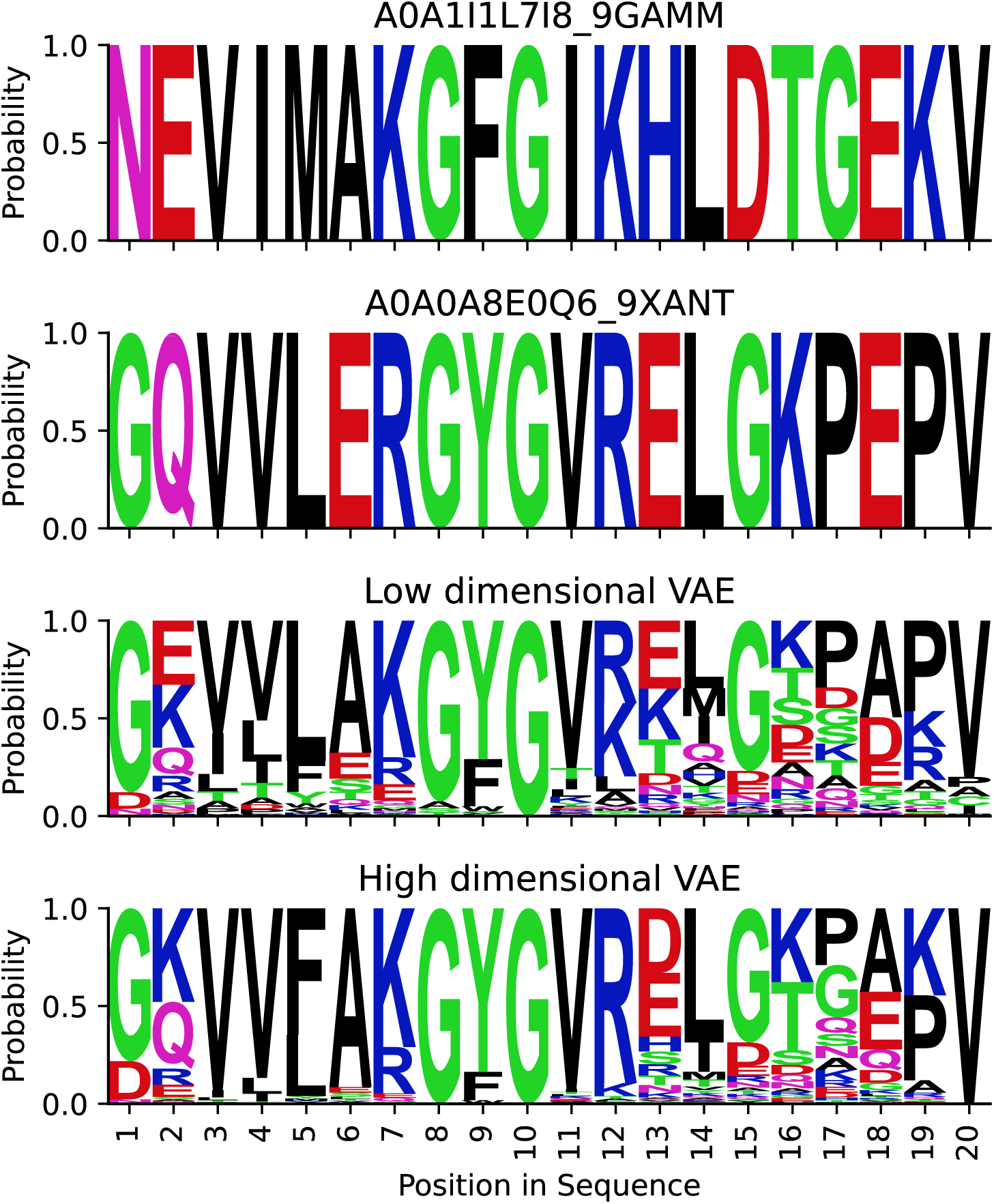
First 20 positions of the sequences A0A1I1L7I8_9GAMM and A0A0A8E0Q6_9XANT and probability logo plot of VAE’s reconstructions of these positions in their parent A35368 (see Figure C2).

## Notes

### Competing Interest Statement

The authors have declared no competing interest.

### Summary of Updates

Major revisions, additional discussion on issue of "blurriness", improved figures.

